# What uniparental genes tell us about the prehistoric human colonization of the Americas

**DOI:** 10.1101/2025.08.04.668419

**Authors:** Vicente M Cabrera

## Abstract

**OBJECTIVES:** From the perspective of uniparental markers, the view of the human prehistoric settlement of America is that it resulted from a single main migration after the Last Glacial Maximum, following a long or short period of genetic isolation in Beringia. Ancient DNA and whole genome analyses have confirmed this view. My objective here is to demonstrate that humans entered America before the LGM and that the documented post-LGM expansions began in South instead North America.

**METHODS:** In this work, I have reanalyzed all publicly available mitochondrial DNA and Y-chromosome haplogroups in the American population using simple phylogenetic and phylogeographic methodologies.

**RESULTS:** The arrival of the American settlers occurred more than 30,000 years ago, preceding the LGM. As genomic studies have uncovered, at least two Asian populations contributed to the ancestry of the immigrant population. Low population density and climatic deterioration led to a long period of demographic eclipse, during which small bands of hunter-gatherers made long journeys in search of favorable niches. After the LGM, the climate improved, and demographic expansions occurred in multiple independent centers. The founding and expansion ages of the uniparental haplogroups indicate that these centers were in South America, particularly the Colombian isthmus, the Andean region, the Southern Cone, and the Amazon. Subsequent dispersals occurred in North America, one involving mitochondrial haplogroups A2, C1, and D1, but not B2, and another, of lesser magnitude, represented by the expansion of haplogroups C4c and X2a.

**DISCUSSION:** This work offers a previously unexplored model for the colonization of the Americas.

**RESEARCH HIGHLIGHTS:** Since the past century, the predominant molecular genetic theory explaining the prehistoric human colonization of the Americas has been that Asian hunter-gatherers entered North America across a Beringia land bridge during or after the LGM. Complementary hypotheses have resolved the evident problem of crossing the existing ice sheets at that epoch, such as the LGM Beringian standstill or the existence of a Pacific maritime coastal route during that period. In contradiction, here, the reanalysis of uniparental genes shows that people were already in America before the LGM and that the post-LGM expansions detected with all kinds of molecular markers occurred first in the southern areas of the continent instead of North America.

## INTRODUCTION

The interpretation of the prehistoric peopling of the Americas has been a multidisciplinary enterprise ^1,2^. From a molecular genetics perspective, the first foundations of these interpretations are based on the information provided by the uniparentally inherited mitochondrial DNA (mtDNA) and the non-recombining portion of the Y-chromosome (NRY) in extant populations ^3,4^. As ancient DNA (aDNA) samples are a mixture of endogenous aDNA and contaminating DNA, results on human samples were first questioned. Fortunately, new sequencing technologies have overcome this problem ^5^, and current aDNA-based analyses have added an invaluable timing perspective to these interpretations ^6^. Later, technological improvements have made it possible to sequence entire genomes in present-day and fossil individuals. Analyses of these genomes have revealed extinct ancestries and genetically well-differentiated ancestries from the past that, admixed in different proportions, persist in current Native American populations along the American continent. This new information has improved previous hypotheses and formulated new ones ^7^.

Thus, some conclusions have achieved general acceptance: 1) The Asiatic origin of the Native American ancestors; 2) The first migration, in one or several waves, occurred as a rapid movement towards the south of the double continent; 3) The initial low population density led to strong genetic differentiation geographically structured; 4) Subsequent migrations reshaped the genetic landscape of North and South America.

Necessary for the last was the entrance from Beringia of several migratory waves that originated the settlement of Paleo-Inuit and Inuit; 5) Time comparisons of population genetic profiles since post-Neolithic to present-day have uncovered an essential genetic continuity (although see **<u>Llamas et al</u>**. ^**8**^).

However, other propositions are still under debate, such as: 1) the precise Asian origin of the Native American ancestors; 2) the routes followed by these pioneers as they expanded throughout the Americas; and, perhaps, the most contentious question, 3) when these first migrations occurred. Although the most recent studies based on complete genomes favor a post-Glacial dispersal ^9^, with or without a prolonged standstill in Beringia ^10^, the first works with mitochondrial DNA favored an early entry, around 30 kya before the Last Glacial Maximum (LGM), avoiding the problem of having to cross the ice sheets of North America ^11,12^. These age discrepancies are due to the molecular evolutionary rates used in each case. Unfortunately, the rate of molecular evolution is not constant; it depends on several parameters, one of which is the adequate population size (Ne), which has changed dramatically from the Paleolithic to present-day ^13^. Therefore, like radiocarbon dating, molecular rates must be calibrated to obtain more accurate ages. One way to do this is by using ancient genomes and/or mitogenomes, but in the case of the American colonization, all the extracted aDNA postdates the LGM ^6^. The option is to use evolutionary rates calibrated with Paleolithic fossils to date Paleolithic events ^14^. Despite their utility, whole-genome studies barely mention uniparental markers ^15^. However, see ^9,16^. The underlying reason for this abandonment is that these markers only offer monogenic insights, and only the set of all genes provides congruent results. In my opinion, this assumption is flawed. The leading cause of genetic differentiation in estimating phylogenies and genetic distances between populations or species is isolation. Isolation affects whole genomes and does not discriminate by sex. Populations differentiate genetically, due to the loss of previous polymorphisms, by fixation or extinction, and the accumulation of new polymorphisms by mutation in a time-dependent way. Thus, in the majority of the cases, any DNA fragment, including those coding for uniparental markers, should be valid to realize genetic comparisons whenever they have accumulated enough genetic differentiation. The generalized use of DNA barcoding in molecular taxonomy ^17^ validates my previous assertion. Isolation is broken by migration/introgression, affecting uniparental markers and autosomes differently. Whether uniparental markers persist only in their original populations or both endure in the original and recipient populations, their coalescence node—the distance to their most recent common ancestor—will remain unchanged. However, if the introgressed fragments are not thoroughly masked for mixed autosomes, they will influence the genetic distances obtained between the populations/species involved. A paradigmatic example is the controversial phylogenetic relationships obtained between modern humans and their closest relatives, Neanderthals and Denisovans. While mtDNA and NRY sequences showed modern humans and Neanderthals as sister lineages ^18,19^, autosomes joined Denisovan and Neanderthal as the closest pair, leaving modern humans as an out-group ^18^. Contrary to the widespread acceptance that the proper relationship between hominids is that obtained with the autosomes, I have suggested that the valid phylogeny is deduced from the uniparental markers. In contrast, the one obtained from the autosomes resulted from later introgressions between the three groups of hominins ^20^. The fact that the mtDNA Primate phylogeny is strongly congruent ^21^ and that the mtDNA and Y-chromosome great ape subspecific phylogenies are totally concordant ^22^, whereas the human genome shows more than 36% of their polymorphisms incongruent with the primate species tree due to incomplete lineage sorting ^23^ are in support of my assertion. Taking into account the above-mentioned particularities of the uniparental markers, in this work, I perform a thorough reanalysis of all published mitogenomes and the most recently published Y-chromosome results, using phylogenetic and phylogeographic tools, to reconsider and shed new light on the hypotheses formulated to explain the prehistoric colonization of the New World by Asian peoples. The main conclusions obtained are: 1) that the first migration involved at least two Asian populations with most probable central and eastern Asian origins; 2) that this wave(s) occurred before 30 Kya; 3) that those pioneer colonizers did not stop in Beringia but kept trekking across the Americas leaving few genetic traces in past and present-day native American populations; 4) That after the LGM, small and relatively isolated resilient groups radiated from different areas to occupy new exploitable territories both in North and South America; 5) that these secondary expansive waves occurred first in southern-an meso-America than in northern North America; and 6) that gene flow existed among geographically adjacent groups.

## RESULTS

All the phylogenetic and phylogeographic mtDNA results described below have been obtained from the trees presented in the supplementary figures (Supplemental_Fig_S1.xls (HgA2), Fig_S2.xls (HGB2), Fig_S3.xls (HgC1), Fig_S4.xls (HgD), and Fig_S5.xls (HgX2), and all the NRY results described below have been obtained from the trees already published by other authors ^24–30^.

### The most probable origin of the American Asian ancestors

The main Native American mtDNA haplogroup founders can be divided into two groups: 1) those belonging to macro-Haplogroup M (C1, C4c, D1, D2a1, D4e1b, and D4h3a), and 2) those that belong to macro-Haplogroup N (A2, B2, and X2a). The same grouping occurs when we look at the number of mutations accumulated in their basal stems that link them to the common nodes with Asian lineages. While C1, C4c, D1, D2a1 and D4h3a have two mutations, and D4e1b only one; A2 and B2 have five mutations and X2a four mutations (**Table 1**).

**Table 1.** Mutations on American basal branches to their common nodes with Asia.

| American haplogroups | Mutations | Coalescent Age (Kya) | Asian haplogroup | Asian region |
| --- | --- | --- | --- | --- |
| A2 | 5 | 20,2 | A-a1 | NW China (Xinjiang) |
| A10a | 1 | 4,04 | A10 | Western Siberia |
| B2 | 5 | 20,2 | B4b1a | Altai |
| C1 | 2 | 8,08 | C4 | Eastern Asia |
| C4c | 2 | 8,08 | C4a'b'c | Eastern Asia |
| D1 | 2 | 8,08 | D4 | Eastern Asia |
| D4e1b | 1 | 4,04 | D4e1 | Eastern Asia |
| D2a1 | 2 | 8,08 | D2a'b | Northeastern<br>Siberia |
| D4h3a | 2 | 8,08 | D4h3 | NW China<br>(Xinjiang) |
| X2a | 4 | 16,16 | X2a'j | Tajikistan |

An unpaired t-test shows that the difference between groups is highly significant (t = 7.2164; df = 5; two-tailed P value = 0.0008). Furthermore, it is well known that macro-Haplogroup M is dominant in southern and eastern Asia ^31^, while macro-Haplogroup N is more frequent in western Eurasia ^32^. Consequently, the same occurs with the Asian ancestral sequences/haplogroups that show the shortest distances with the Native American haplogroups. It has already been demonstrated that the closest relatives of the macro-Haplogroup M American lineages are in eastern Asia/southeastern Siberia ^33,34^. Some further commentaries are pertinent for these lineages: Hg C1 comprises four common and well spread sub lineages, three of them (C1b, C1c, C1d) practicably exclusive of the Americas and one (C1a) with a wide Asian range, and several rare clades/isolates, the most ancient of which is represented by two identical sequences from Brazil ^35^ and Paraguay ^36^ that carries twelve mutations in its basal stem (Supplementary_Fig_S3.xls). The other isolates have been detected in northwestern Eurasia instead of East Asia: C1e in Iceland ^37^, western Russia ^38^, and western China ^39^; C1f in Tajikistan ^40^, and C1g in Mesolithic remains from western Russia ^41^. As the oldest clades are found in the New World, the most parsimonious conclusion is that all the C1 branches in Eurasia resulted from retro migrations from America. The oldest Native American samples harboring C1b lineages are USR1 from Alaska ^42^ and I11974 from Chile ^43^, both around 12,000 years old; for C1c, it is MW291667 from Argentina 41, approximately 8,500 years old; and for C1d, it is Lapa01, 9,600 years old, from Brazil ^43^. Haplogroup D1 has the oldest samples in Yucatan, Mexico, represented by a Late Pleistocene skeleton (HN5/48) around 12 ky old ^44^, in Spirit Cave, Nevada, around 11 kya ^9^, an approximately 10 ky old LLP.S2.E1 sample from Argentina ^45^, and the Sumidouro4, 9,645 years old, sample from Brazil ^9^. Until recently, the American haplogroup D4h3a was separated from its Asian root by six mutations, but recent aDNA studies ^46^ have rescued a Chinese sequence that shares four of the six basal substitutions with the American clade (Supplementary_Fig_S4.xls). The remains, from which this sequence was extracted, were unearthed at the Amur region, in northeastern China, and are around 14,000 years old. The oldest samples harboring D4h3a sequences in the New World are the 12,600-year-old Anzick1 from Montana ^47^, the 10,300-year-old Shuká Káa from Alaska ^48^, and the 10,400-year-old specimens from Lagoa Santa in Brazil ^9^.

About the macro-Haplogroup N, for the New World haplogroup A2, the closest Asian sequences are BZL_M16A and BZL_M16B from haplogroup A-a1 (according to the YFull MTree nomenclature) that belong to historic samples from Xinjiang, western China ^39^. For haplogroup B2, it is AY519494, included in the Asian haplogroup B4b (Supplementary_Fig_S2.xls), with the Altai region as its geographic origin ^49^. Finally, for the American haplogroup X2a, the closest Asian relatives are into haplogroup X2j, that groups samples from Egypt and Europe (Supplementary_Fig_S5.xls) but, for geographic proximity, I consider as the closest Asian sequence a partial one (sample 40) from Tajikistan ^50^, as it carries 16179 and 16357 substitutions that are terminal mutations in the Egyptian X2j ^51^ mitogenomes (Supplementary_Fig_S5.xls). For haplogroups A2 and B2, the most ancient sequences, around 14,000 years old, were extracted from human coprolites found in Paisley Caves, Oregon ^52^, the next oldest for A2 is Lapa15 (around 8,700 years old) from Lapa do Santo, Brazil ^43^, and for B2 is USR2 (approximately 12,000 years old) from Upward Sun River site in Alaska ^42^. The most ancient Native American sample with an X2a haplotype is the Kennewick Man from Washington, which is around 9,000 years old ^53^.

A way to reconcile the close arrivals of the Asian ancestors carrying the Native American M and N lineages with their significant differences in the number of mutations accumulated on their basal stems is to assume that the route followed across Asia of the N carriers was longer and slower than that of the M carriers.

Finally, if we equate the number of lineages carried by the M and N groups to their respective population sizes upon arrival in America, we could estimate that the contribution of the eastern Asian contingent was 62.5% and that of western Eurasia was 37.5%.

The NRY does not show American colonization patterns as clear as those of mtDNA, but using its phylogeny and phylogeography in the Americas and Asia I have considered the ancestors of the American clades C-MPB373, that has its closest haplotypes in the eastern Asian outgroup clade C-L1373 ^26,54^, QF746 that includes the Q-YP1475 subclade represented by the Saqqaq ancient sample ^55^ and has as sister subclade Q-B143 represented by northeastern Siberian Koryaks ^56^, and Q-FT9713, that has its closest sequences in Mongolia and northeastern China (https://www.yfull.com/tree/Q) as of eastern Asian origin, while C-P39/Z30536, having as outgroup the Central Asian subclade C-F1756 ^24^, Q-Z780, that includes the Native American ancient sample Anzick1 ^47^ and has Q-L330 of Central Asia ascription ^57,58^ as sister clade, Q-M3, that has as out group Q-L804 with a northern European geographic range ^59^, and the rare Q-Y1150 and Q-M378 ^25^ as of western Eurasian origin. Unlike mtDNA, most American male lineages appeared to have their ancestors in western Eurasia. Interestingly, the C-MPB373 South American lineage is phylogenetically an outgroup of C-F1699 that comprises numerous western Eurasian lineages, including the Canadian C-P39/Z30536 branch ^26^. Also of interest is that the most abundant American clade, Q-M3 ^28^, has as a sister branch Q-L804, which is specific to northern Europe ^25^. The oldest sample belonging to the eastern group of lineages is Lapa01, from Brazil, which is approximately 8,840 years old ^43^. For the western group, Anzick1 from Montana, with an approximate age of 12,600 years, represents the oldest North American sample ^25^. In contrast, the oldest sample (I11974) in South America was found in Los Rieles, Chile, and is approximately 10,500 years old ^43^.

### The global ages of the first arrival and expansions of the Native American parental ancestors

A global average of the different mean mutational data, calculated for each American mitochondrial haplogroup using various approaches (**Table 2**), give mean mutational values of 7.11 (95% CI: 6.34 – 7.88) substitutions at the arrival in America of the founding lineages, and a mean of 4.42 (95%: 3.98 – 4.85) substitutions for their initial expansions into the American continent, which, in turn, produce mean ages of 28.72 (95%: 25.61 – 31.84) kya and of 17.86 (95% CI: 16.08 – 19.59) kya for the founding and expansion events respectively.

**Table 2.**
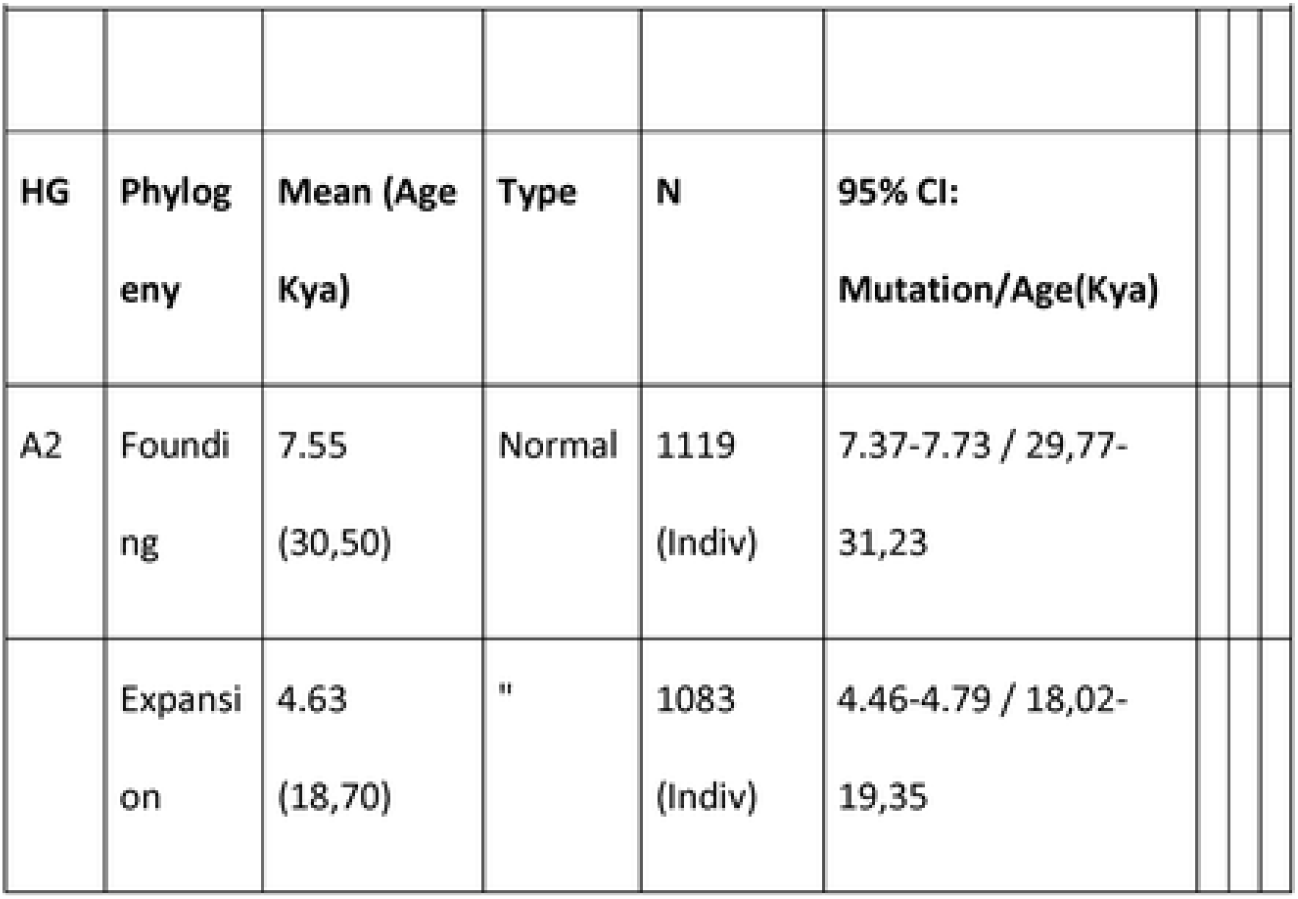
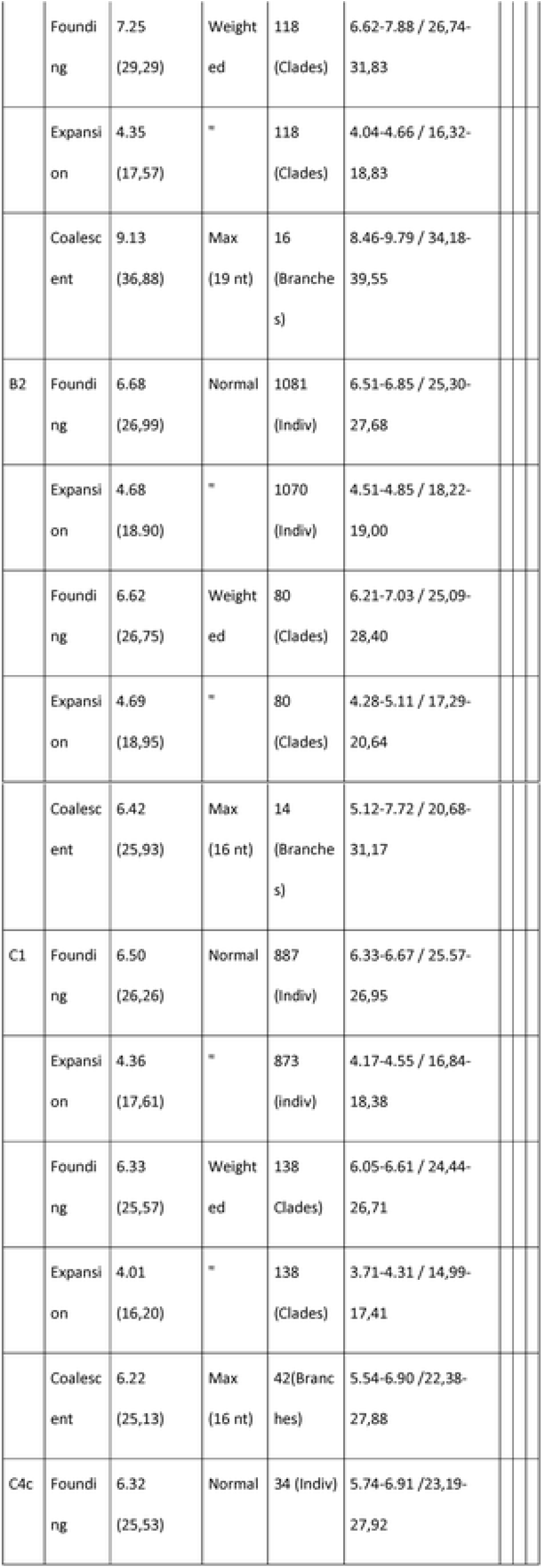
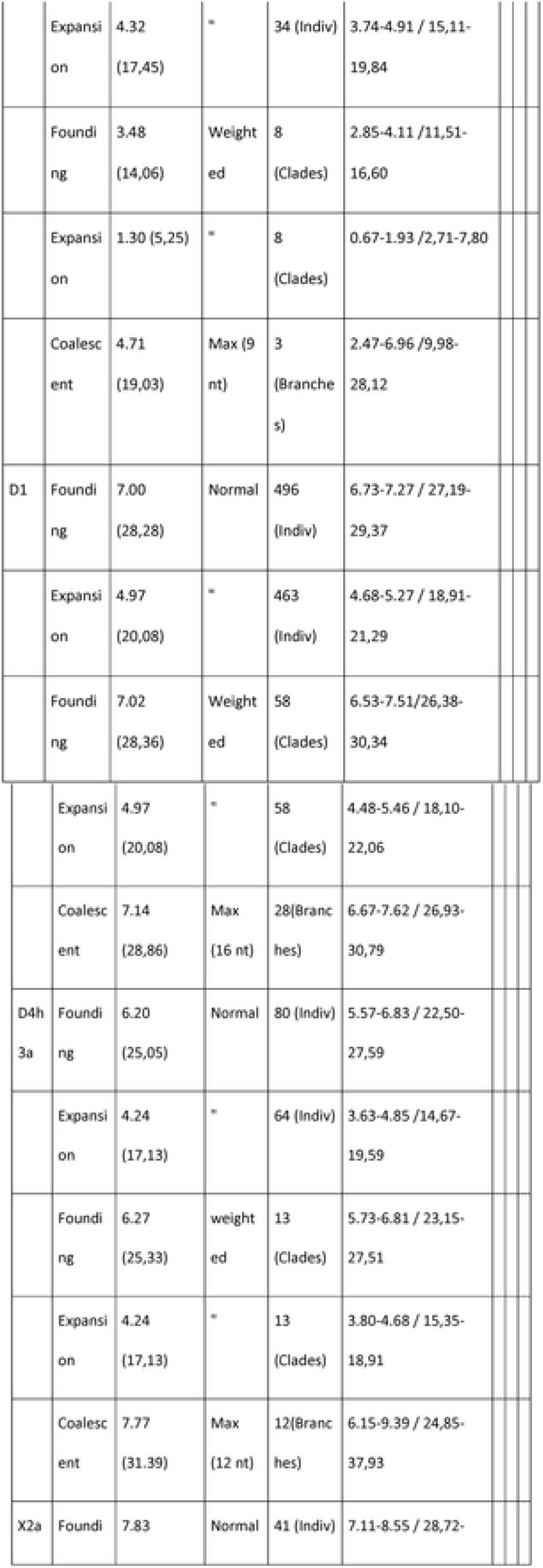
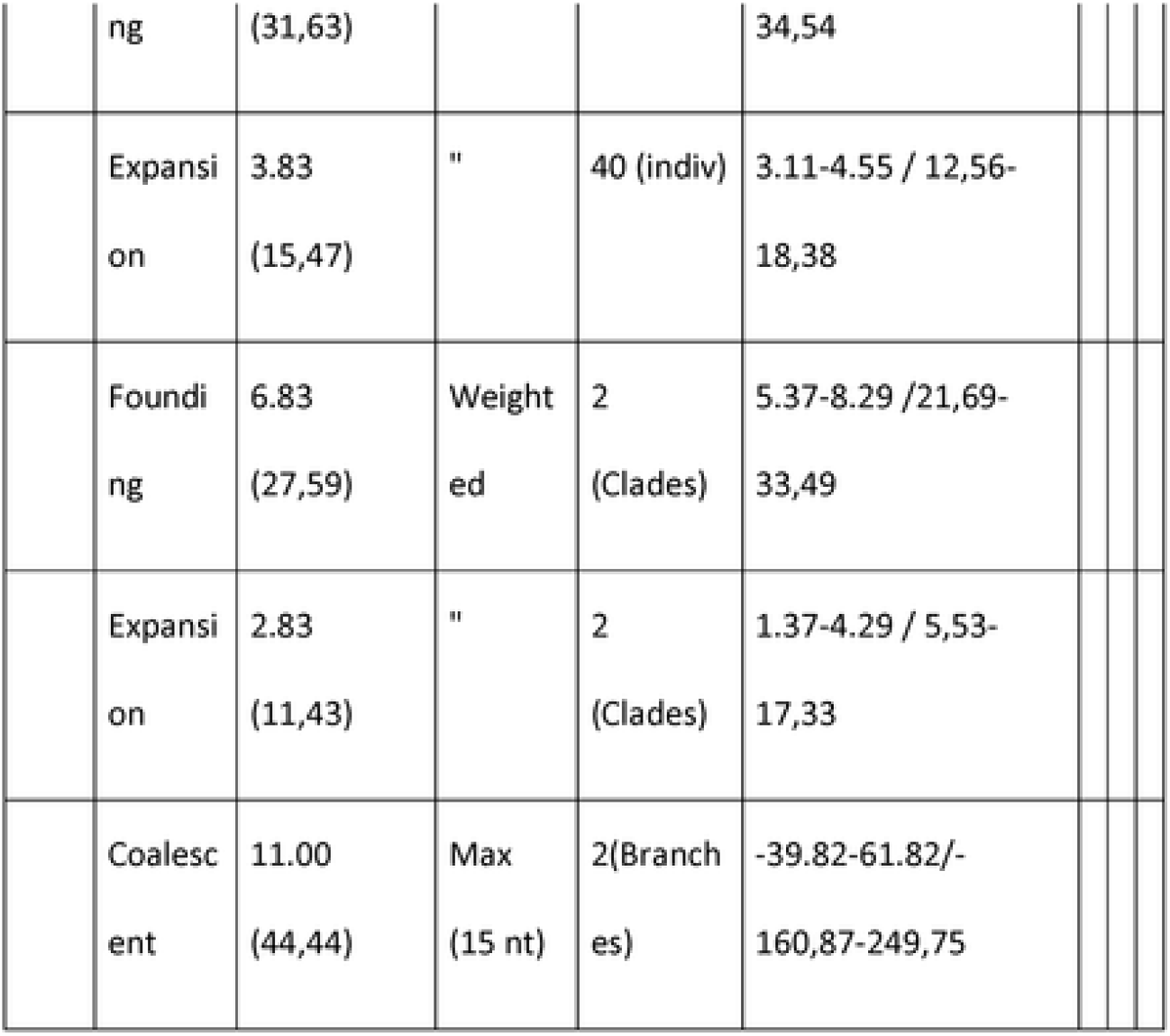
Mutational means and coalescent mean ages for the main American mtDNA haplogroups.

The founding ages for the mtDNA Native American haplogroups, exposed above, coincide with the approximately 30 kya proposed in the first mtDNA studies ^11,12,33^, and with the calibrated ages calculated by **Rieux et al**. ^**60**^, based on ancient mitogenomes. However, recent extant and ancient mitogenome studies ^6,8^ and whole genome-based studies ^7,9^ strongly support post-LGM ages for the settlement of the Americas. However, this is partly inspired by the most conservative archaeological ages proposed for the American settlement.

The age for the first population expansions calculated here, 17.86 kya, after the rigors of the LGM had lessened on the continent, does not differ much from those proposed by most authors. The difference is that this expansion coincides with most authors’ arrival in America proper. In contrast, here, the arrival in America occurred much earlier, around 30 kya, before the glacial period had its worst effects.

Concerning the NRY, it deserves mention that, assuming a first colonization of the Americas about 15 kya, Poznik et al.^58^ estimated a substitution rate for the Y-chromosome of 8.2 × 10-10 (95% CI: 7.2 × 10-10 – 9.2 × 10-10) per site, year. However, doubling the time to 30 kya, as proposed here based on mtDNA, the substitution rate would be 4.1 × 10-10 per site, year which is very similar to that obtained by **<u>Francalacci et al</u>**. ^**61**^ of 5.3 × 10-10 (95% CI: 4.2 X10-10-7.0 × 10-10) per site, year and both estimates falling within the limits of the mean 4.7 × 10-10 (95% CI: 2.1 × 10-10 – 9.5 × 10-10) per site, year proposed by myself based on past population sizes obtained under phylogenetic considerations ^14^. These results contrast with most NRY younger coalescence ages estimated by other authors ^26,28,29,62^.

As YFull Tree v13.03.00 (<u>YTree</u>) has the most actualized NRY phylogeny and the most extensive phylogeography, it is increasingly used as a data source in scientific articles, as in the present work. The YFull substitution rate ^63^ is identical to the one calculated by **<u>Poznik et al</u>**. ^**64**^ under the premise that the human American settlement occurred around 15 kya; therefore, I have halved their substitution rate values when I used them. Under these requirements, NRY haplogroup C clades would arrive in the Americas in separate waves, C-MPB373 first at about 32.2 kya (95% CI: 28.8 – 35.6 kya), and C-P39/Z30536 second at about 26.2 kya (95% CI: 22.8 – 29.8 kya). The three main NRY haplogroup Q clades have very similar ages around 30 kya: Q-F746, 31.8 kya (95% CI: 28.4 – 35.2); Q-Z780, 31.0 kya (95% CI: 29.0 –33.0); Q-M3, 29.8 (27.0 – 32.8). The age of arrival to the Americas of the rare clades Q-M1150 (30.0 kya (95% CI:26.6 – 33.4 kya) and Q-M378, 21.4 kya (95% CI: 17.6 – 25.4 kya), that include European and Asian lineages, is only tentative (https://www.yfull.com/tree/Q).

### Regional heterogeneity for founding and expansion events

Initially, I compared the founding and expansion ages of native haplogroups between the North and South American subcontinents (**Table 3**). Except for haplogroup C1, the results show significantly older ages in South America for both founding and expansion events.

**Table 3.** Differences in Founding(F) and expansion(E) ages between the North America and South America subcontinents.

| Haplogroup | t-value | df | p-value | oldest region |
| --- | --- | --- | --- | --- |
| A2 - F | 3.0397 | 1119 | 0.0024 | South America |
| A2 - E | 4.8326 | 1105 | <0.0001 | South America |
| B2 - F | 1.0604 | 1099 | 0.2892 | ns |
| B2 - E | 3.1395 | 1083 | 0.0017 | South America |
| C1 - F | 1.4777 | 499 | 0.1401 | ns |
| C1- E | 0.4722 | 482 | 0.6370 | ns |
| D1 - F | 5.7700 | 498 | <0.0001 | South America |
| D1 - E | 2.2473 | 466 | 0.0251 | South America |
| D4h3a - F | 3.0245 | 69 | 0.0035 | South America |
| D4h3a - E | 1.0115 | 59 | 0.3159 | ns |

More detailed comparison between geographic regions, within and among subcontinents, using mean molecular age comparisons, pairwise genetic distances and Pearson correlation tests are described in supplementary tables (Supplemental Table S1.1 to S1.7 for A2; Supplemental Table S2.1 to S2.7 for B2; Supplemental Table S3.14, .7, .10, .13, .16 for C1b; Supplemental Table S3.2, .5, .8, .11, .14, .17 for C1c; Supplemental Table S.3.3, .6, .9, .12, .15, .18 for C1d; Supplemental Table S3.19 for C4c; Supplemental Table S4.1 to S4.6 for D1; Supplemental Table S4.7 to S4.9 for D4h3, and Supplemental Table S5.1 to S5.3 for X2a).

### Haplogroup A2

Globally, A2 is the most abundant haplogroup, representing approximately 44% of the Native American population, but it has an uneven geographic distribution. The oldest founding ages for A2 (Supplemental_Table_S1.5) are in Brazil and the Colombia-Isthmus area (≈ 34 kya), expanding first in Colombia around 23 kya. From the Colombian Isthmus region, significant latitudinal clines exist to Siberia (Slope −13.37; R^2^ 0.8396; P-value 0.0102), and to Tierra del Fuego (Slope 23.49; R^2^ 0.8091; P-value 0.0377). In addition, notice that in the most ancient samples of southern South America, mtDNA haplogroup A and B lineages were totally absent or very rare ^45,65–68^ while they were detected in the most ancient samples recovered in northwestern North America, more than 14 kya ^52^, and also were prominent in ancient samples from the Isthmus-Colombian area ^69^. Another geographic peculiarity of haplogroup A2 is its significant abundance in the eastern regions compared to the western ones ^35,70–72^. Notably, the oldest A2 haplotypes are found in North America and Brazil (Table S1.4). Furthermore, a secondary southward migration of some North American sub-lineages is also detectable. One of these lineages is A2f, which is significantly older in North America (≈35,000 ya) than in Mexico (≈23,000 ya) (t=3.6172; df 22; p-value 0.0015). For haplogroup A2, the shortest molecular genetic distances between regions (Table S1.3) were found for the pairs CSAm–SSAm (0.657), NWSAm–NESAm (0.682), and NCAm–NNAm (0.681). When these distances are shown in a PCoA (**Fig. 1**), the first principal coordinate places NESAm in a central position, with the remaining regions of North- and South-America positioned to its left and right, respectively. The second principal coordinate separates the areas of each subcontinent as latitudinal clines, reflecting the Pearson correlation results.

**Fig. 1.**
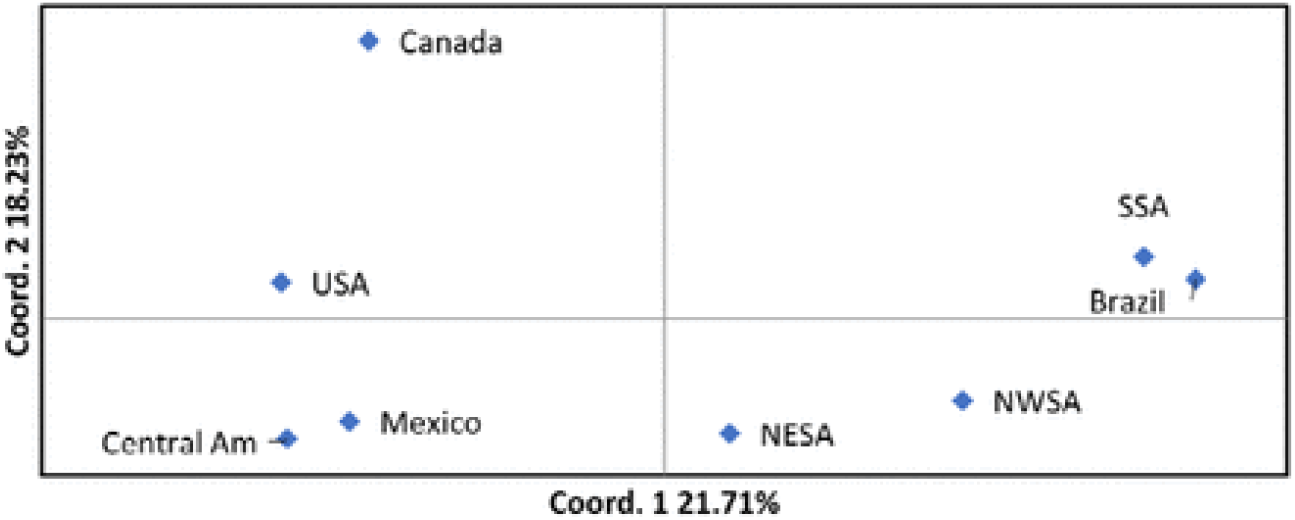
Hg A2 PcoA.

### Haplogroup B2

Haplogroup B2 is the second-largest Native American haplogroup, comprising approximately 24.0% of the global distribution. The oldest founding (approximately 30,000 years ago) and expansion (approximately 21,000 years ago) regional values for this haplogroup are found in South and Central America. As one moves northward from these southern regions, the values decrease in a clinal trend, following a northwest-southeast orientation to reach Central and North America. A significant negative correlation exists between the molecular ages of haplogroup B2 and latitude (Slope: −13.57; R^2^: 0.5455; P-value: 0.0061).

Unlike haplogroup A2, haplogroup B2 is either absent or rare in northern North America, which mirrors its low frequency in northeastern Siberia ^33^. Moreover, in contrast to haplogroups A2 and D1, haplogroup B2 is more prevalent in western continental regions than in eastern ones ^73,74^. The oldest B2 haplotypes have been identified in the Andean region.

The shortest genetic molecular distances were observed between the South South American (SSAm) and Central South American (CSAm) groups (0.492), as well as between the SSAm and Northwest South American (NWSAm) pairs (0.500). A PCoA bidimensional graphic representation of these distances shows NWSAm having a central position, with SSAm and CSAm at its left and NESAm and all northern American regions at its right, following a latitudinal cline as observed with the Pearson correlation (**Fig. 2**). (Supplementary_Table_ S2.3).

**Fig. 2.**
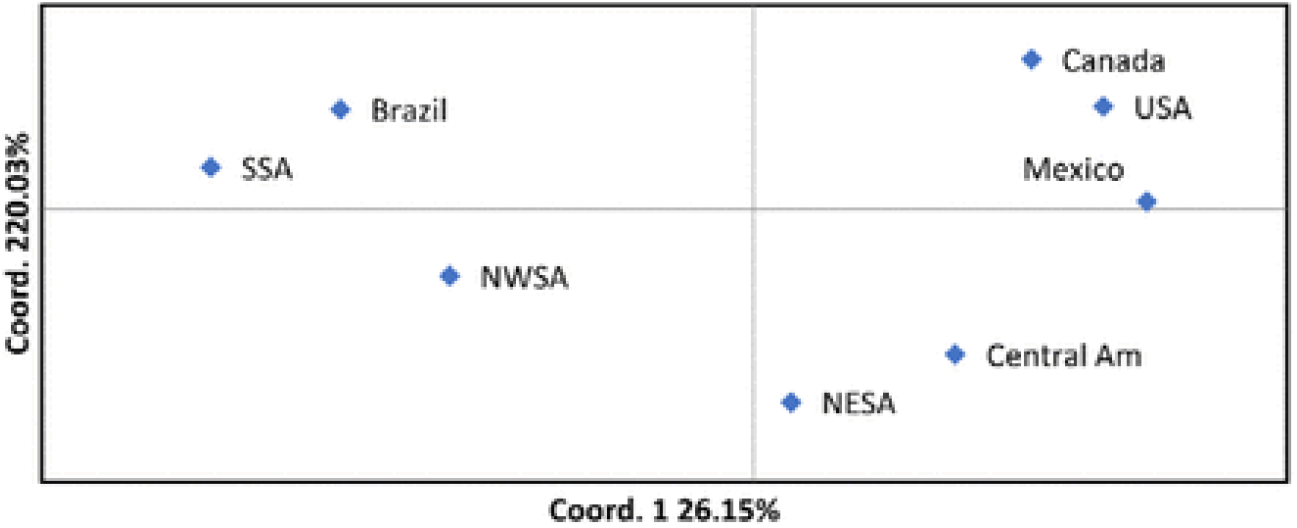
HgB2 PCoA.

### Haplogroup C1

Haplogroup C1 was initially considered a homogenous clade; however, its three main native branches reflect distinct phylogeographic movements that warrant particular attention.

For haplogroup C1b, the oldest founding and expansion regional ages (see Supplementary Table S3.13) are found in Northeast South America (approximately 29,000 to 17,000 years ago), with younger ages extending down to Tierra del Fuego and up to Canada. The data shows a significant trend (Slope −12.20; R^2^ 0.8876; P-value 0.0166). These phylogeographic characteristics are somewhat similar to those of haplogroup A2. However, the longitudinal differentiation seen in the A2 clade is not present in C1b. The oldest haplotypes for C1b have been identified in northern South America, particularly in Ecuador and Colombia, and extend into Mexico (see Supplementary Table S3.10). Additionally, the shortest genetic distance between regions (refer to Supplementary Table S3.7) is observed between Northwest South America (NWSAm) and Northeast South America (NESAm) with a distance of 0.522, followed by NWSAm and Southern South America (SSAm) at 0.605, and SSAm to Central South America (CSAm) at 0.654.

In this context, the PCoA (Principal Coordinates Analysis) graphic representation emphasizes the close relationship between NWSAm–Mexico and the USA–Canada regions, positioning NESAm as the central point from which movements occurred both southward across Brazil and northward into Central America (see **Fig. 3**).

**Fig. 3.**
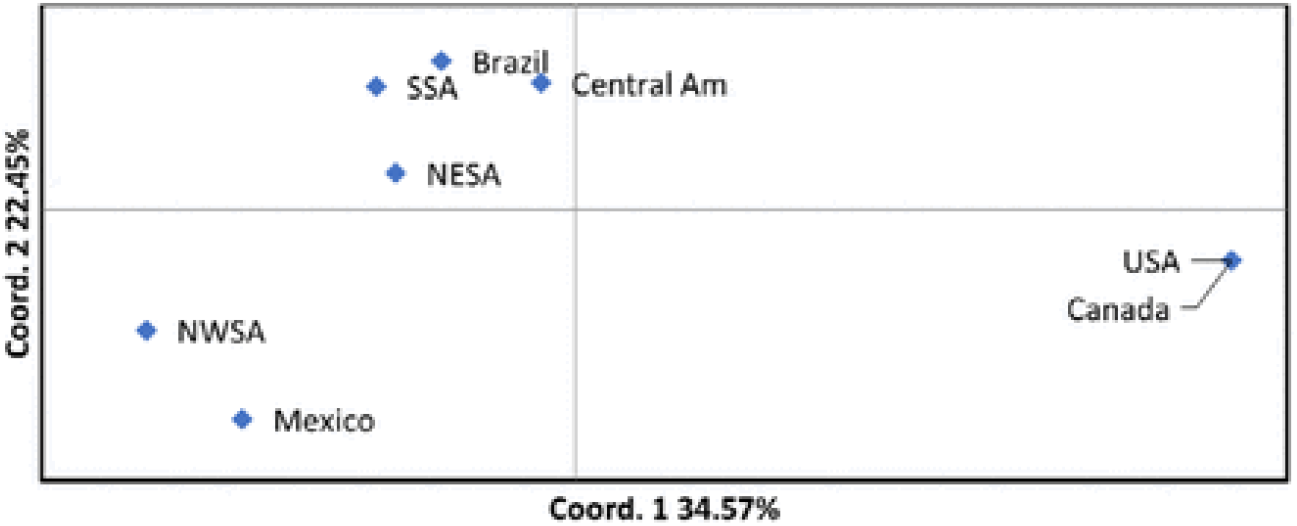
HgC1b PCoA.

For Hg C1c, the earliest regional ages associated with founding and expansion (Supplementary_Table_ S3.14) are centered in NWSAm (≈ 34 kya / 26 kya). From this center, populations with younger ages follow southern and northern geographic transects ending in SSAm and Canada, respectively. The latter shows a significant correlation with latitude for both founding (Slope −14.79; R^2^ 0.8055; P-value 0.0152) and expansion ages (Slope −11.71; R^2^ 0.9629; P-value 0.0005). These results partially resemble those obtained for Hg B2 in its North American subcontinent range. Congruent with its Andean center, the oldest haplotypes for C1c are found in NWSAm and Mexico (Supplementary_Table_ S3.11). The shortest genetic distances within Hg C1c (Supplementary_Table_S3.8) indicate the existence of three geographic centers: the first represented by the CSAm-CAm pair (0.571), the second by the SSAm-NWSam pair (0.630), and the third by the NWSAm-Mexico pair (0.644). This pattern could be explained by supposing NWSAm as the leading center that spread downward to Argentina and Chile and upward to Mexico, and a later center that, from Brazil, reached Central America, erasing the signal of the previous route from NWSAm to Mexico in this region. In this respect, it is worth mentioning that the Y-chromosome Q-M925, mainly Mesoamerican, has close phylogenetic ties with the Brazilian Q-Y26547 subclade ^28^. This complex pattern is partially reflected in the PCoA plot (**Fig. 4**), which shows a latitudinal cline from NWSA to Canada passing through NESAm, with the unusual inclusion of SSAm in it, and an independent close relationship between Central America and Brazil.

**Fig. 4.**
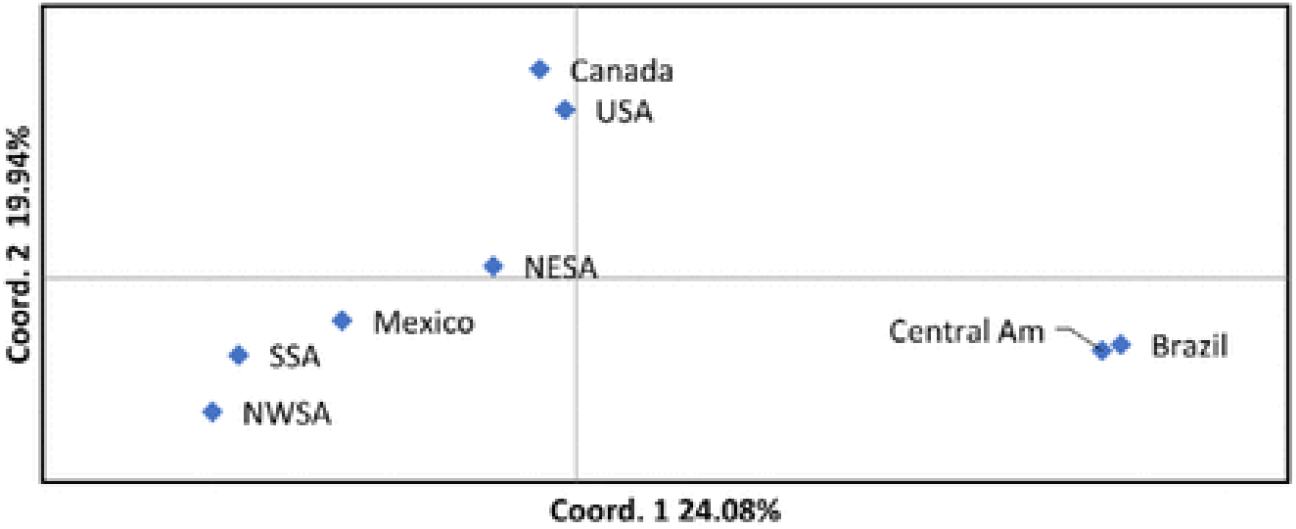
Hg C1c PCoA.

Hg C1d, with the oldest founding and expansion ages (Supplemental_Table_S3.15) in CSAm and SSAm (approximately 38 kya / 33 kya), is the oldest C1 branch. In fact, it is the oldest Native American clade. Although not reaching statistical significance, C1d shows a geographic trend of decreasing population ages from North America to Central America (Supplemental_Table_S3.15). This pattern resembles the one observed for Hg A2. Additionally, it exhibits a significant latitudinal correlation from SSAm to Central America, with the oldest founding (Slope −16.65; R^2^ 0.7649; P-value 0.0415) and expansion ages (Slope −13.29; R^2^ 0.6985; P-value 0.0382) in southern South America. This geographic trend within the South American subcontinent mirrors the pattern seen in Hg B2. The oldest C1d haplotypes are found in CSAm and SSAm (Supplemental_Table_S3.12), supporting the idea of an ancestral southern South American center of expansion for this clade. For C1d, the closest genetic distances (Supplemental_Table_S3.9) are between the SSAm-CSAm pair (0.591) and the NWSAm-NESAm pair (0.592). The latter reflects the close genetic relationship between these regions, also seen in the C1b clade. The PCoA bi-dimensional representation of these distances (**Fig. 5**) shows a central group linking SSAm and CSAm with Central America, and two distant groups: one on the left joining NWSAm and NESAm, and another on the right grouping the northern North American populations and Mexico. The independent and close genetic relationship between the South American regions and Central America resembles what was found for C1c, which was explained by a later expansion from Brazil to Central America. In contrast, the close genetic affinity between NWSAm and NESAm mirrors what was observed in C1b.

**Fig. 5.**
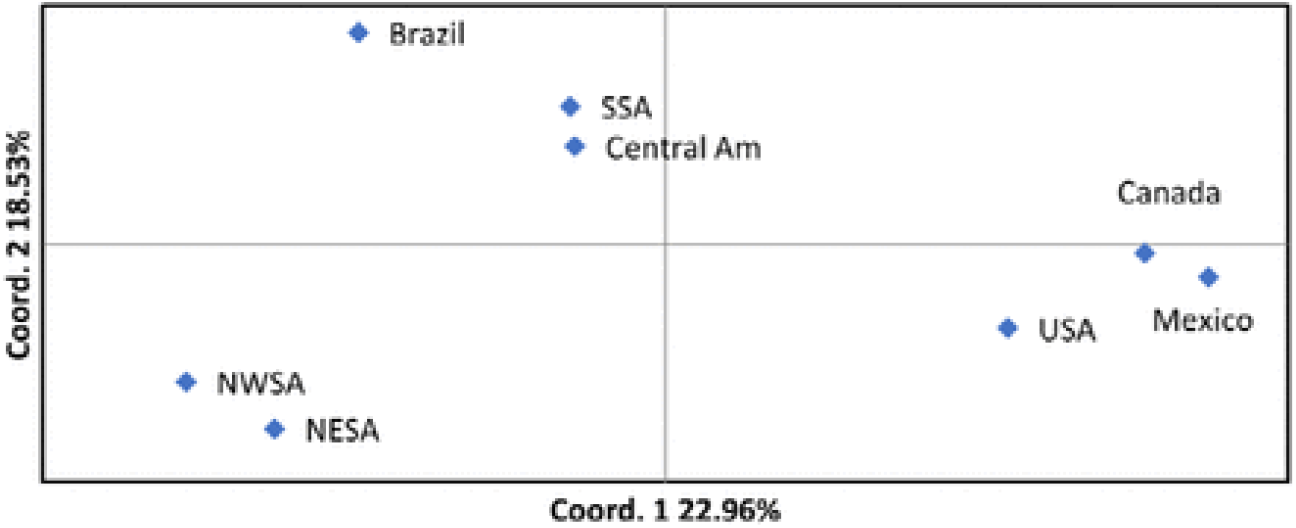
HgC1d PCoA.

### Haplogroup C4c

The founding and expansion ages for C4c are approximately 25 and 17,000 years ago, respectively (**Table 2**), but ages for its main branches vary significantly (Supplemental_Table_S3.19). The most developed branch, C4c1, has founding and expansion ages of about 16 and 12,000 years ago, following the LGM. In contrast, the less common C4c2 sister branch has a founding age of 31 years ago, aligning more closely with other main Native American haplogroups. Still, its expansion age is very recent, at roughly 3,000 years ago. C4c is a minor haplogroup with a limited geographic distribution that, aside from a sporadic find in Colombia and an ancient sample from Brazil, Loca Do Suin_9100 years old ^75^, is confined to NNAm (Supplemental_Fig_S3). Therefore, the unusual results observed for this lineage may result from limited sampling and/or strong drift effects.

### Haplogroup D1

For D1, the oldest founding ages (around 31,000 years ago) are found in SSAm and CSAm (Supplementary_Table_S4.5). The earliest expansion ages are also identified in these regions, but at different times (CSAm approximately 23,000 years ago; SSAm approximately 18,000 years ago). Regarding regional D1 founding ages, there is a significant latitudinal correlation from SSAm to Central America (slope −16.65; R^2^ 0.5431; P-value 0.0370), while a non-significant decreasing trend is observed from Canada to Central America (Supplementary_Table_S4.5). The oldest D1 haplotypes are primarily located in NWSAm and later in SSAm (Supplementary Table S4.4). The shortest genetic distances in D1 are between SSAm-CSAm (0.618), USA-Canada (0.667), and those involving NWSAm with SSAm (0.768) and with NESAm (0.759). A PCoA plot (**Fig. 6**) visually illustrates these genetic relationships, placing all Mesoamerican and northern South American populations in a central position, distinctly separated from two groups—one consisting of northern North American populations and the other of southern South American populations, which are very distant from each other.

**Fig. 6.**
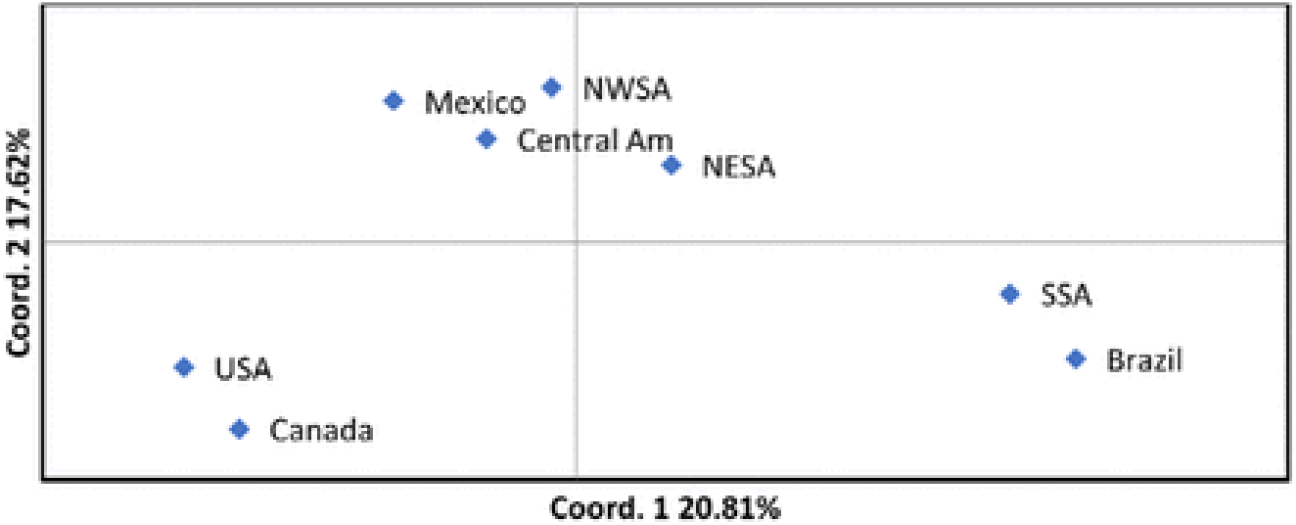
Hg D1 PCOA.

The D1 phylogeographic data described above closely resemble the patterns observed for haplogroups B2 and C1d.

### Haplogroup D4h3a

For D4h3a, the oldest founding age (approximately 30,000 years ago) and expansion age (roughly 22,000 years ago) occurred in CSAm (Supplementary_Table_S4.8). Age comparisons between regions are only significant when the oldest expansion ages of SSAm and CSAm are compared to those of NNAm and NWSAm (Supplementary Table S4.9). When living separately, one can observe that NESAm is isolated; the oldest D4h3a haplotypes are found in NWSAm, Mexico, and consistently in Chile (Supplementary_Table_S4.7). It has been suggested that D4h3a indicates a Pacific southward expansion ^76^. However, these results do not support that idea. It appears that D4h3a was more common in the past and was most frequent in eastern regions compared to today.

### Haplogroup X2

Haplogroup X2a is a minor Native American haplogroup with a limited geographic range confined to northern North America ^76^. The estimated ages for the origin and spread of haplogroup X2a are approximately 29 years ago and 16 years ago, respectively (Supplementary_Table_S5.1), with the oldest haplotypes found in the X2a1 branch in samples from the North American Great Plains. These ages are notably older in North America than in Canada (Supplementary Table S5.3). However, the discovery of a divergent X2 haplotype in Canada ^76^, which may define a new Native American haplogroup called X2g (Supplemental_Fig_S5), is noteworthy. Although the current distribution of X2a, both ancient and modern, is concentrated in the northeastern part of North America ^76^, with sporadic cases in Mexico ^77,78^, its oldest known sample, the Kennewick Man, approximately 9000 years old and carrying a basal X2a haplotype, was excavated in northwestern Washington state ^53^. This finding raises the possibility that the X2a clade may have originated in the northern Pacific region ^76^.

### NRY haplogroups

On a global scale, two major parental lineages with Pan-American distribution have been identified (Q-CTS1780/Z780 and Q-M3), both within haplogroup Q, with the first occurring at lower frequencies everywhere compared to the second. Additionally, two minor lineages belonging to haplogroup C, with a subcontinental range (C-MPB373 and C-P39/Z30536), are considered Native American founder lineages. The southern C-MPB373 has been consistently reported in present-day Colombians and Equatorians ^4,30,79,80^. The northern C-P39/Z30536 has been found in Canada and northern North America, reaching Mexico ^81^, showing a northern to southern gradient of decreasing diversity ^57^. Although phylogenetically Q-CTS1780/Z780 is the oldest clade, the demographic expansion following the LGM mainly occurred among males within the Q-M3 lineage that branched off simultaneously, mainly in Mesoamerica (Q-M925) and South America (Q-M848), forming well-differentiated regional subclades ^25–30,62,70,79,82–84^. Some of these minor Q subclades are also noteworthy due to their geographic distributions and STR diversity. One such lineage is Q-MEH2, identified in northeast Siberian Koryaks. Its close relationship to the extinct Paleo-Inuit Saqqaq ^55^ suggests it could trace a recent migration from Siberia to America across the Bering Strait ^85^. However, Q-MEH2 has recently been found in Nahua populations from southern Mexico, and based on its STR variability, it appears to have arrived much earlier in Mexico than in northern America ^27^. Another notable example is the Mexican Q-PV3 lineage, initially considered a minor branch of Q-L54, but its divergent haplotypic variability suggests it might represent another founding Native American lineage ^27^. This subcontinental differentiation of male ancestries supports diverse demographic histories in North and South America ^82^.

However, before drawing conclusions, it is necessary to examine what NRY ancient DNA analyses indicate, especially since post-Columbian admixture heavily impacted the Native American male genetic pool in both North ^86^ and South America ^87^. The picture was similar. Although Q-CTS1780 might have been more common in the past across both subcontinents ^25,43,75^, Q-M3 was also the dominant NRY lineage in all ancient DNA data examined ^6^. Nonetheless, the detection of today’s extinct lineages in the past ^28,43^ supports the idea that firm founding and drift effects occurred in NRY Native American lineages. These findings do not align well with the hypothesis of a rapid and recent migratory wave from the north to explain the prehistoric colonization of South America.

## DISCUSSION

The early arrival of Native uniparental lineages in the Americas, over 30 kya, as proposed here, renders the hypotheses of a prolonged ^88,89^ or brief ^26^ standstill in Beringia, a previous expansion of their ancestors into Asia ^24,54,90^, or the existence of a Pacific Coastal Route around 20 kya ^91–93^, unnecessary.

The archaeological evidence for humans in the Siberian Arctic around 30 kya ^94^ and possibly since 45 kya ^95^ supports the idea of an early entrance into America by its native ancestors. More challenging is gaining acceptance from archaeologists for human occupation in the American continent before and around the LGM, but it looks pretty likely that this occurred around 16-18 kya at Monte Verde, Chile ^96^, at the Santa Elina site in Central Brazil during the LGM, at Chiquihuite Cave in Central Mexico about 30 kya ^97^, in the Colorado Plateau more than 36 kya ^98^, at White Sands, New Mexico, during a stratigraphic record spanning from 24 kya to 17 kya ^99^, and at Bluefish Caves, Canada, approximately 24 kya ^100^.

Ancient genome studies have characterized the ancestral Native American population as a mixture of northern Eurasians and northern East Asians ^101^. The same dual contribution was previously suggested using NRY data ^4,79,102^, and a similar conclusion has been reached here based on mtDNA data. Genomic research indicates that the ancestral Native Americans arrived in the New World as an already admixed group ^9^. However, although coalescence ages do not show significant differences among haplogroups, the NRY phylogeny is more consistent with two successive waves: the first carrying C-MPB373 and Q-Z780 lineages, and the second comprising carriers of C-P39/Z30536 and Q-M3 lineages. Evidence for mtDNA is less conclusive, but the fact that the basal branches connecting haplogroups C and D to their respective Asian roots are significantly shorter than those of A and B could support the idea that C and D arrived earlier in America. Another clue might be found in the Southern Cone of South America, the region farthest from the continent’s entry point. Ancient and modern samples from this area show that haplogroups C and D were the first and predominant lineages reaching the region, still the most common today ^65–68,103,104^. The dominance of haplogroups C and D has also been observed in Amazonian populations from Brazil ^105^, suggesting an early southward migration. At this respect, it deserves mention that the old age estimated for mtDNA haplogroup D1g has been suggested as support of a pre-Clovis migration to the southern-cone of South America ^106^. Furthermore, mtDNA diversity for haplogroups C1 and D1 increases from north to south, while North American populations exhibit higher diversity levels for haplogroups A2 and B2 ^107^. In North America, haplogroups A, B, and X have expanded across populations in a clinal pattern, whereas there is no evidence of such a spread for haplogroups C and D ^108^. These patterns are consistent with two successive migration waves. However, genetic drift may also explain them, as samples from other regions show the absence of some founder mtDNA haplogroups. For example, the Kuna from Panama lack C and D ^109^, while the Ayoreo from Bolivia show no A or B haplogroups ^110^. Based on the distribution of two rare mtDNA haplogroups, D4h3 and X2a, with the first spreading along the Pacific coast and the second entering via the ice-free corridor, two migratory waves from Beringia have been proposed ^76^. Nevertheless, for D4h3, the authors noted that its rarity in North America and higher frequency and variation in South America require further explanation ^76^.

Another hypothesis, based on genome studies, suggests that after arriving in North America, the native ancestral population split into two distinct branches known as South Native Americans (SNA) and North Native Americans (NNA) ^9^, or their approximate equivalents, Ancestry A (ANC-A) and Ancestry B (ANC-B), respectively ^91^. While the northern ancestry is mainly limited to Canada and eastern North American regions, the southern ancestry is detectable in the oldest sampled genomic sequence, Anzick-1, which is approximately 12700 years old from Montana ^47^, and in the Spirit Cave sample from Nevada, around 10700 years old ^9^, extending southward to Mesoamerica and South America ^9^. Single-parental markers share the geographic range of NNA genomic ancestry, such as mtDNA haplogroups X2a ^111^ and C4c ^112^, as well as NRY lineages C-P39 ^26^, Q-Y4276 ^25^. The situation becomes much more complex when explaining the transcontinental distribution of the southern component of the Native American genome. The initial dual explanation involving southward Pacific coastal expansion—where successive populations could be seen as mixtures of SNA and NNA components in varying proportions ^91^—was proven insufficient ^43^. First, the uniqueness of the northernmost SNA component, represented by the Clovis-associated Anzick-1 sample (SNA1), could not account for the genomic makeup of many South American populations, leading to the need to add a new, dominant non-Clovis SNA2 component, partly represented by Spirit Cave ancestry, to better explain the genomes of many southern populations. Approaching Mesoamerica, the Mexican Mixe population required gene flow from an unsampled population for accurate representation ^9^. Furthermore, several undiscovered genomic ancestries needed to be included to properly model the structure of various South American regional populations. An interesting case is the presence of a divergent ancestry related to Australasian people found in Amazonian tribes such as Surui, Karitiana, and Xavante ^113,114^. This component, called “population Y,” was confirmed in ancient Brazilian samples like Sumidouro-5, approximately 10000 years old ^9^. At least one mtDNA haplotype, likely from haplogroup B—whose southernmost lineage aligns well with the Australasian ancestry—was necessary to accompany this component. However, identifying the ancestral Asian population from which it entered the New World remains challenging. Basic lineages of haplogroup B appeared in eastern Asia around 40,000 years ago, as shown by the Chinese Tianyuan sample ^115^. The Australasian ancestry shared by ancient South American samples and Tianyuan is also notable ^116^, suggesting it could have arrived with populations deeply rooted in eastern Asia with ancient Australasian ties, or with carriers of mtDNA haplogroup B from central Eurasia. The Isthmus-Colombian region also showed genomic complexity, with a Pleistocene ancestry reaching Panamanians and remaining confined to the Isthmus ^69^. Hunter-gatherers from the Colombian Altiplano also carried an unknown basal lineage from the initial South American genomic radiation ^16^. In northwest South America, a previously unrecognized ancestry was distinctive of northern Ecuador and Colombian populations ^117^, while Native Peruvians from the Lake Titicaca area showed a Siberian component also found in ancient North Americans like Anzick-1, Kennewick, and Saqqaq ^48^. The complex ancestries uncovered in South America have led some scholars to caution against oversimplifying the rapid post-glacial colonization model, suggesting that a northward expansion from South America to explain the Pacific migration wave would require a more intricate history ^43^. Others propose that the dispersal pattern of SNA south of the continent’s ice sheets involved complex admixture events among earlier populations ^9^. The uniparental markers analyzed in this study offer an alternative hypothesis, potentially integrating all genomic data from a different perspective.

If we accept that Asian populations gave rise to Native Americans before 30 kya, then the story of human settlement in America can be viewed differently. It is known that since their arrival, the climate has gradually worsened, covering the Arctic and Antarctic regions with ice ^118^, clearing the Amazon and Central American rainforests^119^, and transforming most of the land into forests and dry steppes ^120^. During this tough period, humans had to survive in small, isolated groups and adapt to different glacial refugia, leaving few archaeological traces. When the climate improved, these groups grew in number and expanded into new, now habitable areas. Thus, unlike the rapid southern wave of colonization proposal, the American continent was not empty of humans. The post-LGM colonization was not a single or multiple waves spreading from northern North America to settle Mesoamerica and then South America. Instead, it involved more or less simultaneous radiations from various regions, most of which, based on uniparental markers, are centered in South America. One major expansion center for mtDNA is the northwest Colombian-Isthmus area, where younger haplogroup A2 and C1b clines are seen stretching north and south. A similar distribution of A2 frequencies has also been observed in this region ^121^. Another potential core is in the Andean region, where mtDNA haplogroup C1c shows its oldest origins and expansion ages, with younger clines extending toward Canada and down to the SSAm. This pattern mirrors haplogroup B, which expanded from Chile and the Andean region into Mesoamerica. However, it appears that at least part of the B2 cline results from displacement by other mtDNA lineages in North America, as B was present early on in northeast Asia as an ancestral B lineage ^115^ and in Alaska as a basal B2 lineage ^42^, before nearly disappearing from both areas. The highest frequencies and genetic diversity of haplogroup B2 are found in the Andean Altiplano ^122,123^, the Bolivian Piedmont ^124^, and the Argentinian Gran Chaco ^104^. Although the classic explanation for the B2 cline suggests descent from California ^123^, South American origins have also been proposed, from the Peruvian Andes ^125^ or the Chilean Southern Cone ^126^. Similarly, the C1d haplogroup, like B2, shows its most ancient origins and expansion in the southern Cone, spreading northward with younger ages toward Central America, indicating a significant geographic pattern. In North America, some lineages of C1d show an expansion history that ends in Central America, paralleling trends observed in haplogroup A2 sub-lineages. C1a ^89^, a northeast Asiatic branch of C1, along with minor European sub-clades C1e ^37^ and C1f ^41^, suggests a later reverse migration from America ^89^. Using the evolutionary rate proposed here, the founding age of C1a in Asia is about 26.02 kya (95% CI: 18.58-33.49 kya), signaling a return to Asia during worsening weather conditions. Its spread across Asia around 13.9 kya (95% CI: 6.46-21.37 kya) points to a Mesolithic migration, similar to the pattern seen in the northeast European C1f clade ^41^. This early American retro-migration might also be supported by the phylogeny and phylogeography of the Y-chromosome haplogroups C-L1373 ^26,28^ and Q-L54 ^25,26^.

Another mtDNA haplogroup indicating the southern Cone region as a key center of expansion is D1, which shows a significant negative correlation from this area to Central America. Its mean coalescent ages are younger as you move upward, although there is a longitudinal bias leaning toward central and eastern South American regions. D1 is most common in the eastern parts of the Andean region, with ancient regional clades in Central Argentina ^127^ and the southern Cone ^128^. Similarly, for A2 and C1d, a southward expansion from Canada to Central America is observed for D1. These expansion patterns might explain the B2 retraction seen in North America. The evidence here does not support the idea that mtDNA haplogroup D4h3a signals a Pacific coastal route for peopling of the Americas (81), later backed by genomic studies (9). The earliest ages for the founding and expansion of D4h3a are found in SSAm and CSAm, and are significantly older than those in NWSAm and NNAm (Supplementary_Table_S4.9_ S4.9). Additionally, although D4h3a is absent today, it has been found in ancient samples from Alaska, Shuká-Káa, 10.3 kya ^48^, from British Columbia, Canada, 939, 6.5 kya ^129^, and Montana, Anzick 1, 12.6 kya ^47^. It was also detected in ancient remains from the Brazilian Amazon, Lagoa Santa, 10 kya ^9^. This pattern closely resembles that of B2. Therefore, rather than modeling a southward Pacific Coastal expansion, both markers seem to be remnants of a broader past occupation displaced by more recent human migrations.

A specific area in the Amazon Basin (CSAm) deserves special mention. It participated in all the mtDNA radiation centers discussed earlier and is known as a significant reservoir of deep mtDNA lineages ^130,131^. It stands out for the ancient presence of very deep-rooted Y-chromosome lineages such as C-L1373* ^28^ and for detecting lineages previously thought to be unique to the northern subcontinent, like Q-Y4303* ^25^. This genetic diversity is also supported by whole-genome studies on ancient specimens ^9^ and current populations ^132,133^. This genetic landscape aligns with the region’s linguistic richness. Notably, out of 71 identified linguistic isolates across the continent, over 45% are in Brazil, compared to 15% in North and Mesoamerica combined ^134^. Future archaeological findings will likely uncover human settlements much older than currently recognized. Furthermore, the differences in genetic and linguistic diversity between the northern (NNA) and southern (SNA) American branches make the idea of simultaneous colonization of both subcontinents from a northward expansion unlikely.

## MATERIAL AND METHODS

### Material

For the mtDNA phylogenetic and phylogeographic analyses, I searched for partial and complete mitogenomes in the NCBI GenBank, www.ncbi.nlm.nih.gov/genbank/; Mitomap, www.mitomap.org/MITOMAP; EMPOP, https://empop.online; Ian Logan 2023, www.ianlogan.co.uk/sequences_by_group/haplogroup_select.htm; Genome Warehouse (GWH) public repository, https://bigd.big.ac.cn/gwh; and AmtDB, http://www.amtdb.org. Additionally, for phylogeographic analyses only, I also searched YFull and MTree. The NRY phylogenetic and phylogeographic analyses were based on the ISOGG database, http://www.isogg.org/tree/; YFull YTree v13.03.00; YTree; and data compiled by other authors^24–29^. For brevity, the Y-chromosome lineages in the text are referred to by the letter of their basic haplogroup and their terminal mutation.

### Phylogeny

Maximum parsimony intraspecific phylogenetic trees ^135^ were constructed manually, and haplogroup nomenclature followed the PhyloTree database Building 17, http://www.phylotree.org. Haplogroup classification was confirmed using HaploGrep 3 3.2.1, https://haplogrep.i-med.ac.at ^136^. Newly identified subgroups not classified in PhyloTree were provisionally named by the first letter and digit of their primary haplogroup, followed by an asterisk and consecutive numbers. Both labels are shown on the respective tree when a more formal nomenclature is available in YFullMTree.

### Evolutionary rates

It is well known that the mtDNA evolutionary rate experiences a time-dependent effect, slowing down as we go back in time ^137^. The main reason for this effect has been linked to differences in population size, with the smallest populations during Paleolithic times and exponential growth in recent times ^138^. To account for this anomaly, I used an estimated Paleolithic mitogenome substitution rate of 1.5 × 10^-8 (95% CI: 0.42 - 9.07 × 10^-8) per site per year (assuming a mtDNA genome length of 16,500 base pairs), meaning one substitution occurs roughly every 4,040 years ^14^, to estimate haplogroup coalescent ages for the early prehistoric settlement of the Americas. Using a similar approach, a proposed mean evolutionary rate of 0.47 × 10^-9 (95% CI: 0.21 – 0.95 × 10^-9) per site per year was used to calculate coalescent ages of the Y-chromosome ^14^.

### Ages

I have used three approaches to estimate haplogroup coalescent ages: 1) applying statistics rho ^139^ and Sigma ^140^ on individuals within haplogroups, along with the evolutionary rates mentioned above; 2) utilizing subclades within main haplogroups to calculate a pooled mean, then applying normal averaging and standard error statistics for mutation rates across haplogroups; 3) since the sample sizes of these clades vary significantly, I calculated weighted averages and weighted standard errors to apply with the same normal statistics; 4) a common discussion in genetics is that haplotypes at low frequency might become much more common within a population, although this is generally an exception; typically, rare haplotypes—those at the tails of a distribution—are likely to go extinct due to genetic drift, while haplotypes near the mean tend to increase in frequency. Given that the Native American founding haplogroups have experienced genetic drift over time, I used the most divergent lineages within them as proxies for their past means to estimate coalescence ages ^14,138^. Finally, I have considered founding haplogroup ages those that include the mutations present in their basal stems and expansion haplogroup ages those calculated from their subsequent branches.

### Phylogeography

For phylogeographic purposes, the American continent has been divided into seven geographic regions: 1) North Northern America (NNAm), including Alaska and Canada; 2) North America, covering the entire United States (Nam); 3) Mesoamerica, comprising Mexico, Belize, Guatemala, El Salvador, Honduras, and Nicaragua (Mam); 4) Northeastern South America or Isthmus-Colombian area, including Costa Rica, Panama, and Colombia; 5) Northwestern South America (NWSAm), also known as the Andean region, including Ecuador, Peru, and Bolivia; 6) Central South America (CSAm), including Brazil, Paraguay, and Uruguay; 7) Southern South America (SSAm), including Argentina and Brazil. Additionally, within these regions, distinctions are sometimes made between inland and coastal Pacific and Atlantic areas and between Highland Andes and Amazonian zones. I have implemented some straightforward algorithms described below to support the phylogenetic and phylogeographic analyses.

### Genetic distances

In addition to the molecular distance based on the number of mutations between two homologous DNA sequences (FST), I have implemented a phylogeographic distance (DXY) based on genetic affinity (IXY) between regions X and Y. This distance is measured by the frequency with which they form sister branches within a haplogroup tree (nXY) relative to how often each region participates as a sister branch within the measured haplogroup (nX and nY).

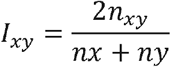

Then D_XY_ is calculated as 1 – I_XY_.

When searching for sister branches between X and Y regions within a haplogroup, I observed cases where one sequence from one of the involved regions served as a direct outgroup to the pair, indicating it as an ancestral relative to the other region, which was considered derived. The significance of this relationship was tested using observed-expected chi-square statistics.

Additionally, for phylogeographic purposes, I have examined each major Native American founder haplogroup to identify the regions where their oldest sub-haplogroups are found and the areas from which they initially expanded.

Finally, to identify the Asian centers where the ancestors of the major American founder clades may have originated, I searched for the most ancient Asian sequences (less mutated) from the Asian-American shared phylogenetic nodes and the regions from which these sequences come.

### Other statistics

Molecular variance (AMOVA), correlation/linear regression, and graphical Principal Coordinates Analysis (PCoA) were analyzed using the GenAlEx 6.5 software ^141^. Descriptive normal statistics, 95% confidence intervals for means, and two-tailed t-tests were conducted with GraphPad https://www.graphpad.com/quickcalcs/. Weighted means and standard errors were calculated using the Weighted calculator (https://calculatodo.com).

### False positive correction

Since I performed multiple statistical tests for each founder haplogroup, I applied a Bonferroni p-value adjustment. However, because the Bonferroni correction is very conservative, I also used the Benjamin and Hochberg ^142^ false discovery rate (FDR) method as a less restrictive alternative.

## Supplementary information

Figures S1 Haplogroup A2 and A10 phylogenetic trees

Figures S2 Haplogroup B2 and B4b1 phylogenetic trees

Figures S3 Haplogroup C1a, C1b, C1c, C1d, C1e, C1f, and C4c phylogenetic trees

Figures S4 Haplogroup D1 and D4h phylogenetic trees

Figures S5 Haplogroup X2 phylogenetic tree

Tables S1 Haplogroup A2 statistical comparisons

Tables S2 Haplogroup B2 statistical comparisons

Tables S3 Haplogroup C1b, C1c, C1d, and C4c statistical comparisons

Tables S4 Haplogroup D1 and D4h3 statistical comparisons

Tables S5 Haplogroup X2a statistical comparisons

## Acknowledgements

Not applicable

